# REDigest: a Python GUI for *In-Silico* Restriction Digestion Analysis of Genes or Complete Genome Sequences

**DOI:** 10.1101/2021.11.09.467873

**Authors:** Abhijeet Singh

## Abstract

Restriction fragment length polymorphism (RFLP) is a technology for the molecular characterization of DNA and widely used genome mapping, medical genetics, molecular microbiology and forensics etc. Terminal restriction fragment length polymorphism (T-RFLP), a variant of RFLP is extensively used in environmental microbiology for the microbial community profiling based on the restriction digestion profile of marker gene (16S rRNA, FTHFS *etc*.) amplicons. At present, there is a lack of a tool which can perform *in-silico* restriction digestion of a large number of sequences at a time, in an interactive way and as an output produce sequences of the restriction fragments and visualization plot. I have developed a graphical user interface based software “REDigest” for the *in-silico* restriction digestion analysis for gene or genome sequences. The REDigest software program with a graphical user interface is freely available at https://github.com/abhijeetsingh1704/REDigest.

## INTRODUCTION

Restriction fragment length polymorphism (RFLP) is a molecular technique used to identify and visualize the variation in DNA sequences. This technique exploits the restriction enzyme target sites in DNA sequences to produce fragments of different length which can be visualized using agarose gel electrophoresis. It is a fast and relatively inexpensive method for the screening and comparing a large number of samples based on their restriction profiles. RFLP or similar techniques have been used in different applications, such as marker-assisted selection (MAS) in animal breeding (Soller, 1978; Beuzen *et al*., 2000), human genetics (Botstein *et al*., 1980), parenthood testing (Jeffreys *et al*., 1985), MAS in plant breeding (Bernatzky and Tanksley, 1986). During the years several variations of this techniques have been developed *i*.*e*. Amplified Fragment Length Polymorphism (AFLP) (Vos *et al*., 1995) and Terminal restriction fragment length polymorphism (T-RFLP) (Liu *et al*., 1997). Especially T-RFLP is being used for the microbial community profiling in environmental and medical microbiology (Osborn *et al*., 2000; Ricke *et al*., 2005; Zhang *et al*., 2008). There are a few programs available for the *in-silico* analysis of restriction fragments like TRiFLe (Junier *et al*., 2008), *in-silico* T-RFLP (Bikandi *et al*., 2004) and NEBcutter (Vincze *et al*., 2003) etc. However, they are limited in terms of output in the form of just numerical values of the terminal restriction fragment size or are low-throughput and can process only a single sequence at a time. During this *in-silico* digestion, only the restriction site position or a restriction map is generated as an output and not sequence information (in any form) is generated. Therefore, there is a need for a program which can process large sequence datasets and generate restriction profile of the whole dataset and still preserves the sequence information in output. Here, I present REDigest, a graphical user interface (GUI) based software program for the *in-silico* restriction digestion of a large number of gene sequences or complete genomes. The name of this software program “REDigest”, is derived from Restriction Enzyme Digestion.

REDigest takes sequence data of genes or a genome in a Fasta (Wikipedia, 2020b; Lipman and Pearson, 1985) or Genbank file format (NCBI, 2006) and produces 1) a Fasta/Genbank format file of the terminal restriction fragment, 2) .csv format file/files (Wikipedia, 2020a) of the restriction and terminal restriction fragments and 3) a restriction fragment diversity plot. REDigest is developed for and can be used on Windows and Linux Operating systems, and tested on Windows 10 (Microsoft, 2015) and Ubuntu Linux derivative Peppermint (Peppermint LLC, 2010). REDigest is licensed under MIT license and can be freely available to download and use from https://github.com/abhijeetsingh1704/REDigest.

## IMPLEMENTATION

REDigest is written in Python3 (The PSF, 2020) (version 3.8.5) and uses module Biopython (Cock *et al*., 2009) (version 1.77) (Chang *et al*., 2020) for the restriction digestion and sequence manipulation, Pandas (The PDT, 2020) (version 1.1.1) for numerical data analysis and Matplotlib (Hunter, 2007) (version 3.3.1) for visualization. The graphical user interface is developed using the Python module Tkinter (Tcl/Tk, 2019) (version 8.6). To use REDigest, it is a prerequisite that all these dependencies must be installed and are present in the execution path.

## USER INTERFACE AND FUNCTIONALITY

### Restriction Digestion and Visualization

REDigest has a very simple and interactive user interface and allow restriction digestion plus visualization in a single program execution from Restriction & Visualization tab. REDigest can process Fasta and Genbank format files as input and can write output file for sequence information in Fasta or Genbank format. The restriction analysis can be performed on a set of multiple genes or a single genome. In case the input file is a multifasta gene file, there is an option to indicate the labelled primer. If the forward primer is labelled, the output will be the sense strand of the terminal restriction fragment and if the reverse primer is chosen as the labelled primer, the output will be the reverse complement sequence of the terminal restriction fragment in 5’ to 3’ orientation. Primer sequences for the forward and reverse primers can also be given which will be added and processed accordingly in the *in-silico* restriction digestion analysis. These options are optional for multigene sequence file. However, these options can be avoided in the case of processing the genome sequence.

The output generated from the REDigest analysis will be different based on the input files. If the input is the multifasta gene file, the output will be 1) a sequence file containing the terminal restriction fragment sequences with unique accession numbers, 2) a scatter plot representing the diversity and abundance of terminal restriction fragments and 3) a data-table containing the information of terminal restriction fragments. This data-table contains the information of the length of terminal restriction fragment and position of consecutive restriction enzyme cutting site. If a whole genome sequence is processed, the output from REDigest will be 1) a sequence file containing the sequences of an individual restriction fragment, 2) a scatter plot representing the diversity and abundance of restriction fragments and 3) two data-tables containing the length of individual restriction fragments and length of terminal restriction fragments of the genome. The interactive GUI of REDigest is presented in **figure 1A**. Users are required to provide the path to the input file and the case-sensitive name of the restriction enzyme, which are mandatory parameters to start the process, indicated in red colour. Rest of the parameters are optional, and the user needs to click the respective button for it to be activated. If the parameters are not provided or if they are not activated, default values will be used during the processing of the input file. The information about all the default values can be accessed from the help menu for respective tabs. Activation of the option will change the label colour to green and the click button will be changed to Ok (**figure 1B)**. On the successful execution of the *in-silico* restriction digestion and visualization process, a pop-up message box will appear on the screen (**figure 2B)**.

**Figure 1:**
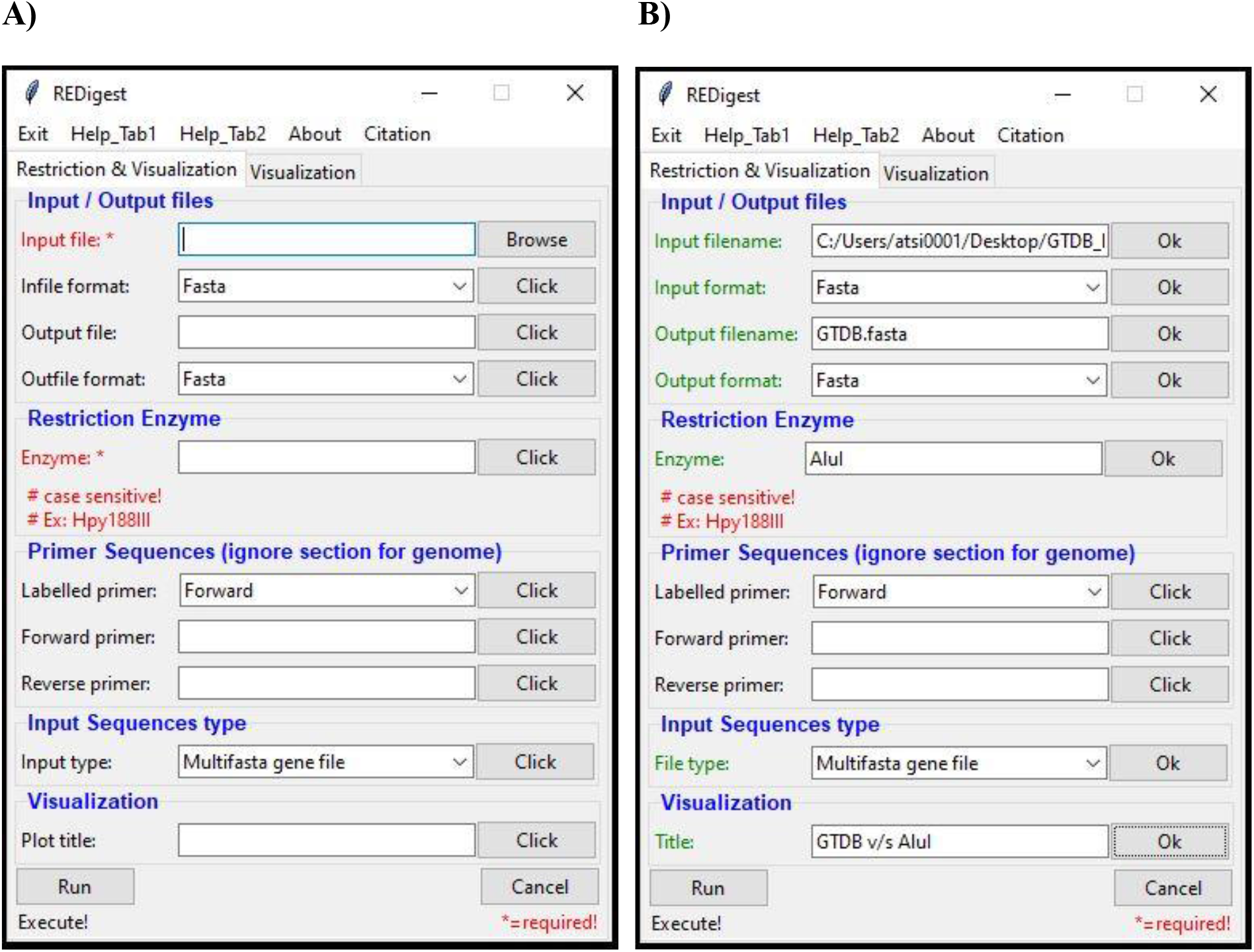
Screen-shot images representing the interactive graphical user interface (GUI) of REDigest software program on the Windows 10 operating system. **A)** the interface upon calling/starting the software GUI. Black colour input labels indicate the value is empty and need to be provided, word required in red colour indicates the respective mandatory field. If the labels (except input file and Enzyme name) are not activated, default values will be used. **B)** User-specific parameters for the respective parameters and its activation is indicated by the change in the colour of labels from black to green. The activation of the parameters is done by the click button which will change to Ok button upon activation.

### Custom Visualization

User can also perform custom visualization from the Visualization tab for the data-table files generated by REDigest from Restriction & Visualization tab. In this visualization process, it is a prerequisite that the data structure in the .csv files is not changed. For the custom visualization, all the plot attributes can be customized as shown in **figure 2A**. After the successful generation of the custom plot, a pop-up message will indicate the completion of the plotting process **figure 2B**.

**Figure 2:**
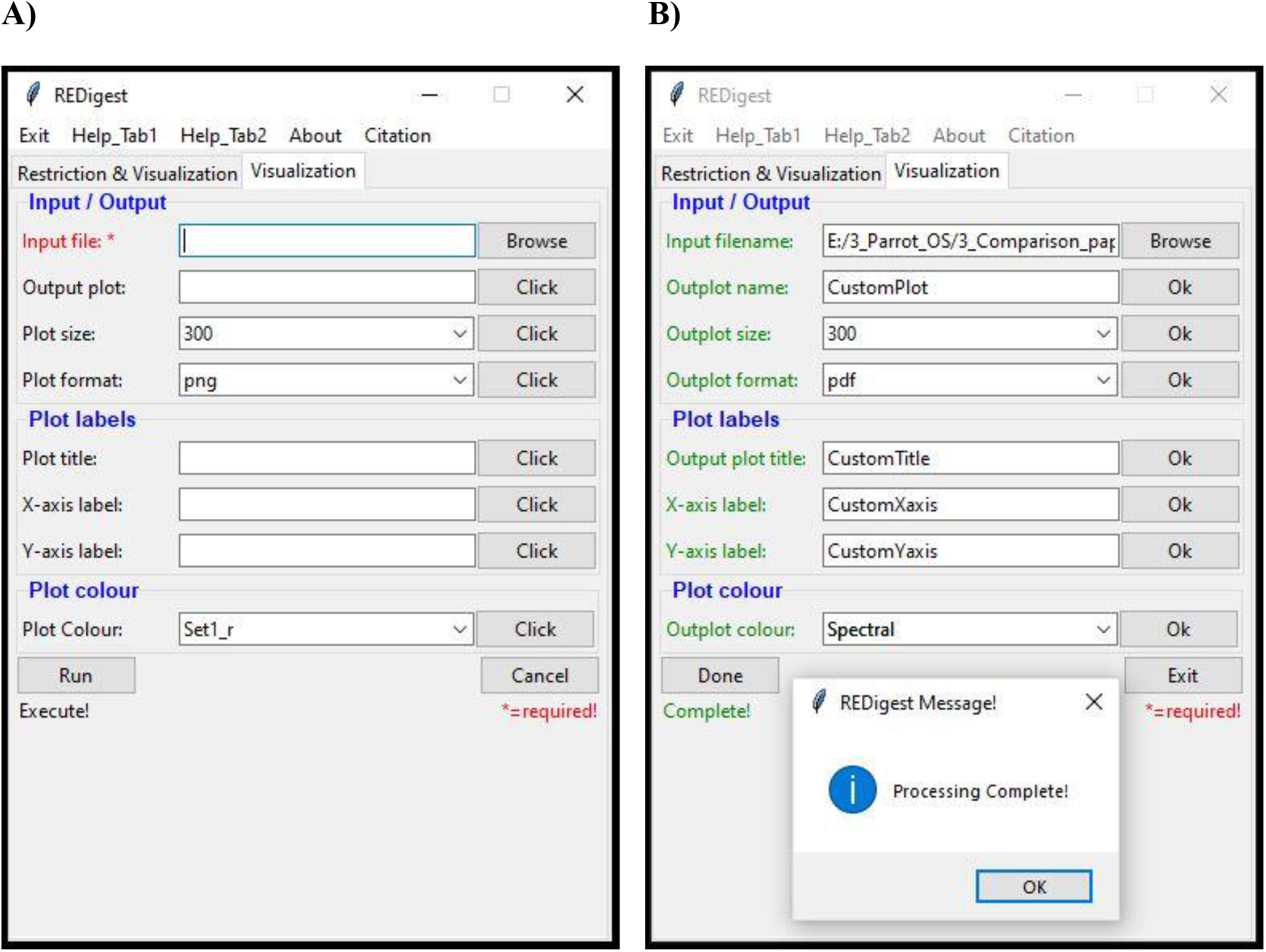
Screen-shot images representing the interactive graphical user interface (GUI) of REDigest software program for the Visualization tab on the Windows 10 operating system.**A)** the window for the Visualization tab where user can provide the custom plot attributes. The input file must be provided by the user, or an error message will appear on the screen. If the respective parameters are empty or not activated, default values will be used. **B)** Custom parameters provided by the user and its activation will be indicated by the respective change of text, colour and click button. A pop-up message will appear on the screen upon successful execution of the process.

### USAGE EXAMPLES

REDigest have been tested against two data-sets of 16S rRNA gene sequences and FTHFS gene sequences from Silva-database training data formatted for DADA2 (McLaren, 2020) and AcetoBase (Singh *et al*., 2019), respectively. 16S rRNA gene training dataset (against restriction enzyme AluI) and AcetoBase dataset contain 374 222 and 6 806 sequences, respectively. These two datasets (Silva and AcetoBase) were processed in REDigest without any marker primer sequence and without any addition of primer sequences with restriction enzyme AluI and Hpy188III, respectively. For the genome analysis, the genome of *Pelotomaculum thermopropionicum* SI (NC_009454.1) was retrieved from NCBI RefSeq assembly and processed against the restriction enzyme AluI. All the analyses were performed on a Windows 10 operating system with 3.4 GHz Inter® Core™ i7-6700 processor and 29 Gb of available RAM. For these two datasets, the output files generated were the same as described earlier for the multigene fasta file. The plot generated from the respective analysis is presented in **figure 3**.

**Figure 3:**
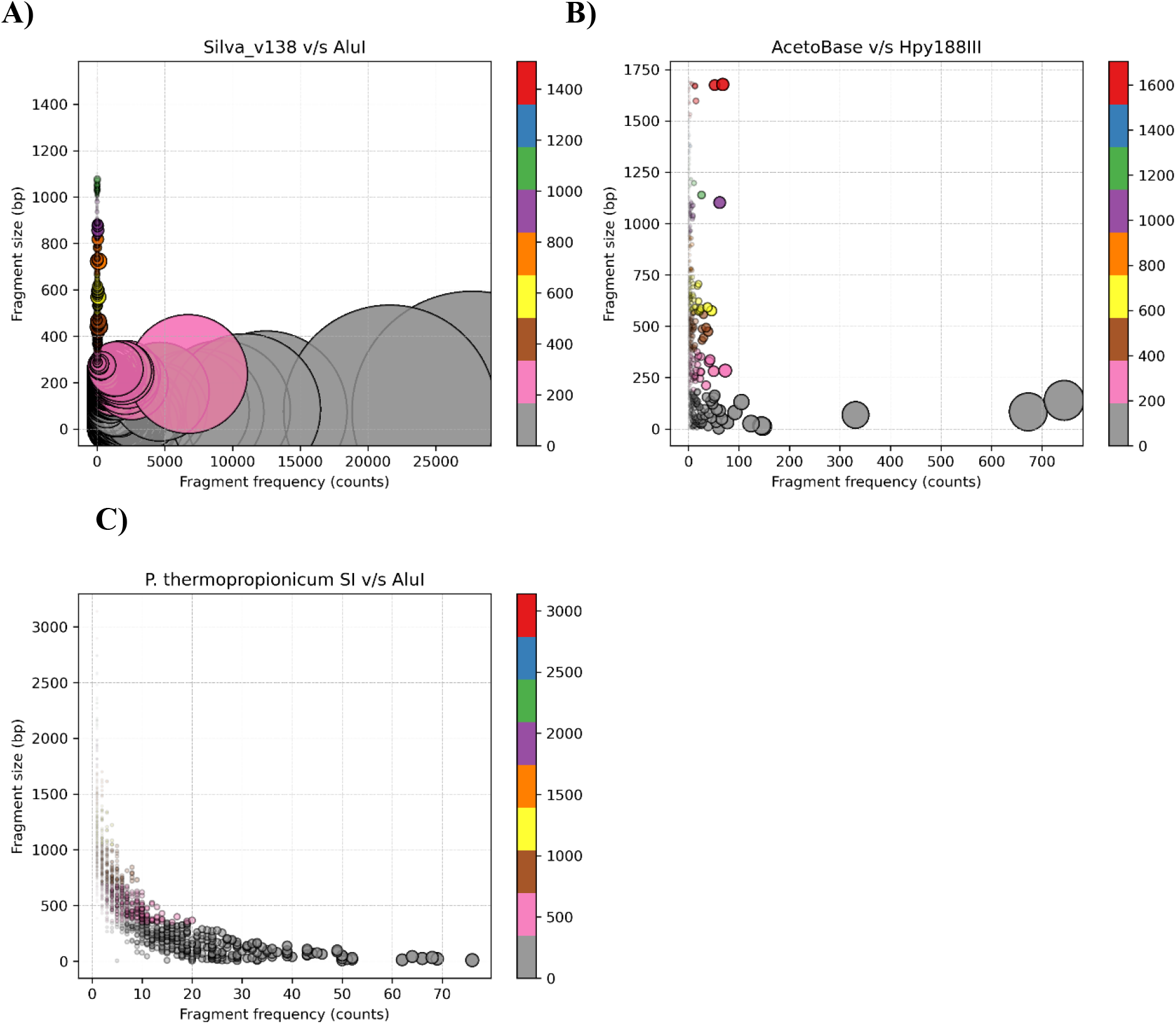
The scatter plot generated from REDigest for **A)** Silva database training dataset processed for the restriction enzyme AluI. **B)** AcetoBase nucleotide dataset processed for the restriction enzyme Hpy188III. C) *Pelotomaculum thermopropionicum* SI complete genome sequence (NC_009454.1) processed against the restriction enzyme AluI.

## CONCLUSION

REDigest is a fast, user-interactive and customizable software program which can perform *in-silico* restriction digestion analysis on a multifasta gene or a complete genome sequence file. REDigest can be helpful in different ways, for example:

1. It can help in the selection of suitable restriction enzyme for the restriction digestion experiment, by processing respective reference gene/genome data with several restriction enzymes and evaluate and choose the best restriction enzyme.
2. Validation of the restriction fragment or the terminal restriction fragment size and taxonomy against a database.
3. A restriction fragment or a terminal restriction database can be generated by using REDigest which can be helpful in the comparison and validation of the experimental restriction digestion profiles

## ACKNOWLEDGEMENTS

I would like to thank Prof. Anna Schnürer for her support and motivation in the development of REDigest and valuable comments on the manuscript.

## COMPETING INTEREST STATEMENT

None to be declared

## Notes

### Competing Interest Statement

The authors have declared no competing interest.

